# A novel *SERPINE1-FOSB* fusion gene in a case of pseudomyogenic hemangioendothelioma results in activation of intact FOSB and the PI3K-AKT-mTOR signaling pathway and responsiveness to sirolimus

**DOI:** 10.1101/2021.05.22.445285

**Authors:** Jun Ren, Xiaohui Wang, Yulin Zhou, Xin Yue, Shouhui Chen, Xin Ding, Xiaoyong Jiang, Xiaokun Liu, Qiwei Guo

## Abstract

Pseudomyogenic hemangioendothelioma (PHE) is an extremely rare disease that affects mainly the young and more men than women. PHE are multicentric, locally aggressive, have low metastatic potential, and affect multiple tissue planes. Genetic aberrations are frequently detected in PHE and may play important roles in the occurrence, development, and treatment of this disease. In this study, we report a case of PHE with a novel *SERPINE1-FOSB* fusion gene. The fusion introduced a strong promoter near the coding region of *FOSB*, resulting in overexpression of intact FOSB. Immunohistochemical analysis showed overexpression of pAKT and mTOR in tumor cells, suggesting activation of the PI3K-AKT-mTOR signaling pathway. The patient responded well to targeted therapy with sirolimus, an mTOR inhibitor. Our study correlated dysregulation of a specific signaling pathway and the effectiveness of a targeted therapy to a specific genetic aberration. This information may be useful for future investigations of targeted therapeutics and provide a potential predictive biomarker for therapeutic effectiveness in cases of PHE with this genetic aberration.

## Introduction

Pseudomyogenic hemangioendothelioma (PHE) is an extremely rare disease that was included in the 2013 WHO classification of soft tissue and bone tumors.^1^ This disease occurs mainly in the young population and affects more men than women. PHE are typically multicentric, locally aggressive, have low metastatic potential, and affect multiple tissue planes. ^1, 2^

Histologically, PHE is characterized by loose fascicles of spindled and epithelioid cells, abundant eosinophilic cytoplasm, and vesicular nuclei with distinct nucleoli. In some cases, neutrophilic inflammatory infiltrates have been noted.^1, 2^ Immunohistochemically, tumor cells are positive for factor VIII, FLI-1, INI-1, vimentin, MDM2, CDK4, CD31, cytokeratin AE1/AE3, EMA, and P63 and negative for CD34, S-100, SMA, desmin, MyoD1, and HMB45.^2, 3^ Genetically, chromosomal rearrangements that result in the fusion of all or part of the *FOSB* coding region with the promoters of other genes, such as *SERPINE1, ACTB, WWTR1*, and *CLTC*, have been frequently identified in PHE, suggesting that activation of *FOSB* plays an important role in the pathogenesis.^2, 4-11^

In terms of PHE treatment, the multifocal nature and high risk of relapse limit the efficacy of surgical excision.^3, 12^ Conventional chemotherapy has been administered in some cases, with variable efficacy. However, intolerable toxic effects have been noted.^12^ In contrast, several targeted drugs, such as sirolimus, everolimus, and telatinib, have shown efficacy with fewer side effects than chemotherapy.^12-16^ Unfortunately, the low incidence of PHE does not allow investigators to perform systematic clinical trials to evaluate the efficacy of a specific treatment, and long-term follow-up data for a specific treatment are limited. Therefore, no systemic therapies have been officially approved for the management of PHE. Thus, in depth knowledge regarding the pathogenesis and treatment and follow-up data for each case of PHE are valuable for understanding and managing this rare disease.

Here, we report the diagnosis, genetic examination, treatment, and follow-up of a patient with PHE and discuss the correlation of genetic aberrations with signaling pathway dysregulation and therapeutic effectiveness.

## Materials and methods

### Histological and immunohistochemical analysis

Resected tumor samples were fixed in 10% buffered formalin and embedded in paraffin to prepare 4-mm sections. For histological analysis, the sections were stained with hematoxylin and eosin. Immunohistochemical analysis of cytokeratin AE1/AE3, CD31, ERG, INI-1, BCL-2, Ki67, CD34, D2-40, S-100, SMA, and STAT6 was performed using the BenchMark XT automated slide preparation system (Roche). Immunohistochemical analysis of FOSB, pAKT, and mTOR was performed as follows. The paraffin was removed from the sectioned tumor samples with xylene, and the sections were rehydrated in a graded series of ethanol, followed by heat-induced epitope retrieval in a pressure cooker and blocking of endogenous peroxidase activity. Next, the sections were incubated with primary antibodies overnight at 4 °C and then with a rabbit secondary antibody and finally reacted with 3, 3′-diaminobenzidine. The samples were counterstained with hematoxylin. Finally, the slides were analyzed using an E200MV microscope (Nikon).

### Detection of the *SERPINE1-FOSB* fusion gene

RNA was extracted from a freshly resected tumor sample using the RNeasy Mini Kit (Qiagen) and reverse transcribed using the RevertAid First Strand cDNA Synthesis Kit (Thermo Fisher) according to the manufacturer’s protocol. The cDNA was amplified in a 20-µL reaction containing 1× FastPfu Buffer (TransGen Biotech, Beijing, China), 1 U of TransStart® FastPfu DNA polymerase (TransGen Biotech), 2.5 mmol/L Mg^2+^, 0.2 mmol/L of each deoxy-nucleoside triphosphate, and 0.2 µmol/L each forward and reverse primer. Based on a previous study,^4^ an upstream primer targeting *SERPINE1* (5′-AGAGCGCTGTCAAGAAGACC-3′) and a downstream primer targeting *FOSB* (5′-GTTCCCGGCATGTCGTAG-3′) were used. PCR was performed on a ProFlex PCR System (Thermo Fisher) under the following cycling conditions: 95 °C for 3 min, followed by 30 cycles at 95 °C for 30 s, 58 °C for 30 s, and 72 °C for 3 min, with a final extension for 5 min at 72 °C.

The PCR products were examined by electrophoresis on a 1% agarose gel containing 1× GoldView™ Dye (SBSGENE, Shanghai, China) for staining. The recovered PCR products were examined using commercial Sanger sequencing (Sangon, Shanghai, China).

### Ethics statement

Signed informed consent was obtained from the patient for the use of his materials and publication of de-identified clinical data, including photographs. The study was approved by the Research Ethics Committee of Zhongshan Hospital Xiamen University.

## Results

### Clinical diagnosis of the case

A 61-year-old man with no specific history of disease presented with rapidly progressive multifocal nodules and localized pain in the left hip and left iliac region for more than 2 months. The skin lesions were discontinuous, raised, or subcutaneous, 0.3–1.5 cm in diameter, with a smooth surface (Fig. 1A). The physical examination was otherwise normal, and superficial lymph nodes were not enlarged. Magnetic resonance imaging (MRI) revealed multiple lesions in the ilium (Fig. 1C), gluteus medius, gluteus mininus, and subcutaneous region of the left hip (Fig. 1H).

**Figure 1.**
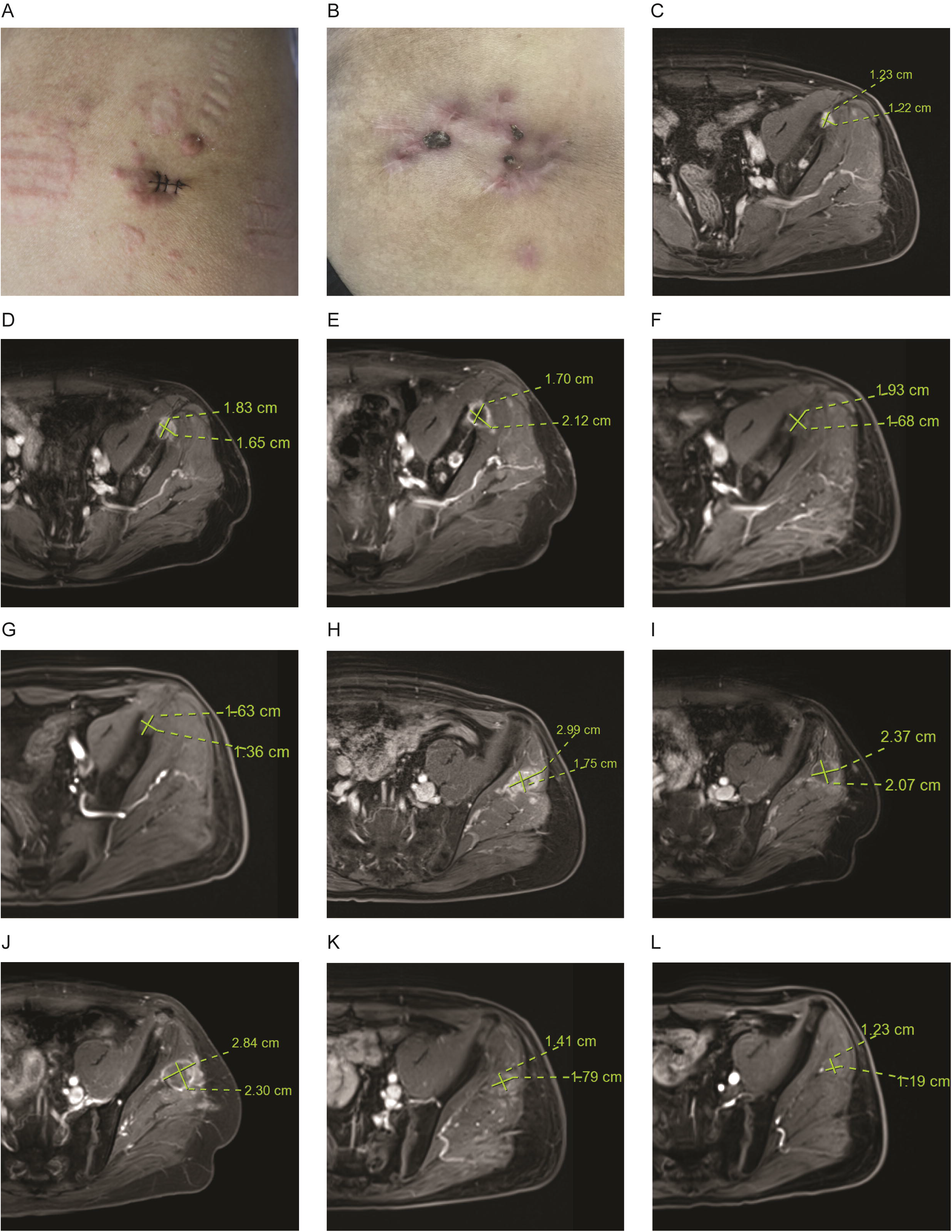
Skin lesions and magnetic resonance imaging of the case. (A) Multiple nodules were present at diagnosis. (B) Remission of the lesions was evident after sirolimus therapy for 2 months. Lesions with high signal intensity in the ilium at diagnosis (C), after chemotherapy (D), after radiotherapy (E), and after sirolimus therapy for 5 (F) and 20 months (G), respectively. Lesions with high signal intensity in the gluteus medius, gluteus mininus, and subcutaneous region of the left hip at diagnosis (H), after chemotherapy (I), after radiotherapy (J), and after sirolimus therapy for 5 (K) and 20 months (L), respectively. The size of the largest lesion was indicated.

As shown in Fig. 2, histopathological examination of a biopsy sample from the left hip revealed that the nodule was in the dermis and subcutaneous tissue and was composed of loose fascicles of spindled and epithelioid cells, with abundant eosinophilic cytoplasm and vesicular nuclei with distinct nucleoli. Neutrophil infiltration and intravascular tumor thrombus were also observed.

**Figure 2.**
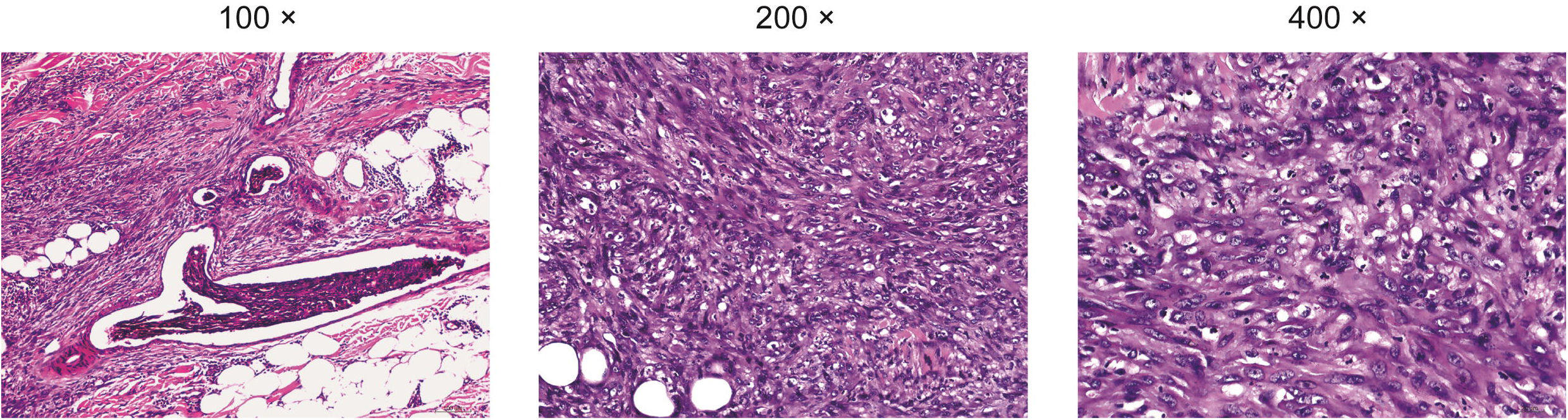
Histopathological examination of a biopsy sample from the left hip.

Immunohistochemical analysis revealed that the biopsy was positive for cytokeratin AE1/AE3, CD31, ERG, INI-1, BCL-2, and Ki67 (15%) (Fig. 3) and negative for CD34, D2-40, S100, SMA, and STAT6 (data not shown).

**Figure 3.**
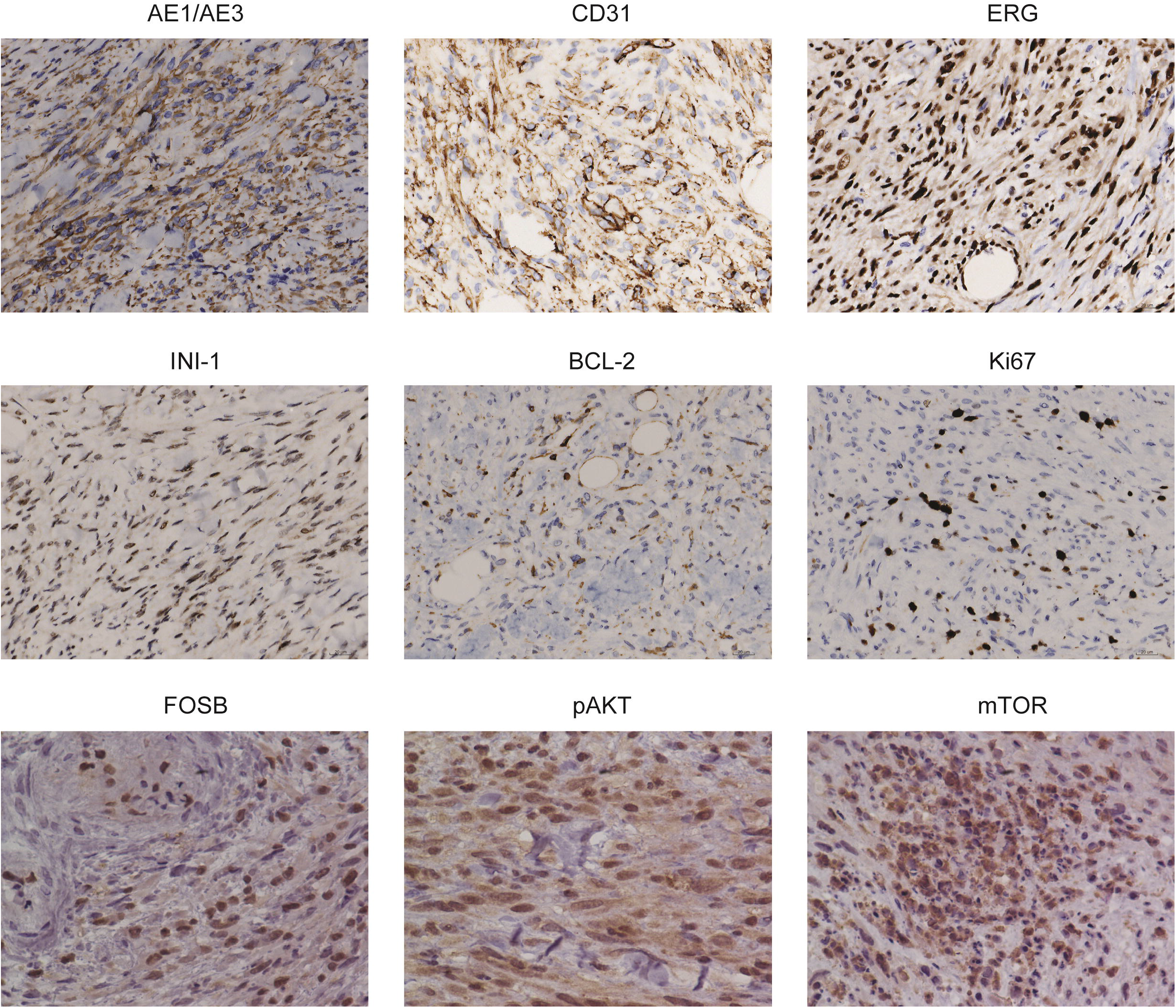
Positive immunohistochemical staining of a biopsy sample from the left hip (400×).

Based on the clinical presentation, histomorphology, and expression of keratins and endothelial markers in the immunohistochemical analysis, the nodules were diagnosed as PHE.

### The tumor was responsive to sirolimus

After diagnosis, the patient underwent surgical resection to remove the palpable nodules and was administered two cycles of anlotinib hydrochloride (10 mg/d) every 2 weeks. During the first cycle of chemotherapy, fresh nodules developed, and pain remained. During the second cycle of chemotherapy, no evidence of tumor development was observed, and the pain was alleviated. However, no evident improvement was noted on follow-up MRI, and chemotherapy was halted due to drug-induced nosebleeds and periungual sclerosis (Fig. 1D and I). The patient was then treated with radiotherapy 5 days a week (Monday–Friday) via 6□MV linear accelerator at a dose of 5000 cGy. Unfortunately, no improvement was noted after 5 weeks of radiotherapy (Fig. 1E and J).

Eventually, targeted therapy with sirolimus, an orally available mTOR inhibitor, was started (1 mg/day), and oxycodone (50 mg bid) was administered for pain control. At 2-month follow-up, no fresh nodules had developed, and the pain was alleviated (Fig. 1B). Moreover, no drug-induced toxic effects were noted on physical examination. Therefore, the dose of sirolimus was increased to 2 mg/day. Follow-up MRI was performed at 5 and 20 months after sirolimus treatment, which showed general improvement of the disease (Fig. 1F, G, K and L), and the largest lesion in the left gluteus medius had progressively shrunk. Since the pain was alleviated, a reduced dose of oxycodone (10 mg each night) was sufficient for pain control. Based on the clinical improvement and lack of side effects, sirolimus therapy and follow-up are ongoing.

### The *SERPINE1-FOSB* fusion gene activates *FOSB* and the EGFR-pAKT-mTOR pathway

Since the nodules were diagnosed as PHE and they were responsive to sirolimus, we investigated the possible pathogenetic and therapeutic mechanisms. We first examined the nodules for the presence of a *SERPINE1-FOSB* fusion gene. As shown in Fig. 4A, when the tumor DNA sample was amplified with primers designed for the presence of a *SERPINE1-FOSB* fusion gene, an amplicon was generated. Sanger sequencing of the amplicon revealed a novel *SERPINE1-FOSB* fusion gene, in which the breakpoints of both genes were in non-coding exon 1. Moreover, a 52-bp fragment from intron 1 of *SERPINE1* was inserted at the fusion junction (Fig. 4B). Since the coding region of *FOSB* was unaffected, the fusion gene could generate an intact FOSB protein. Next, we examined the expression levels of FOSB, pAKT, and mTOR in the tumor. As shown in Fig. 3, strong expression was noted for these targets, suggesting that the fusion gene activates FOSB and the PI3K-AKT-mTOR signaling pathway.

**Figure 4.**
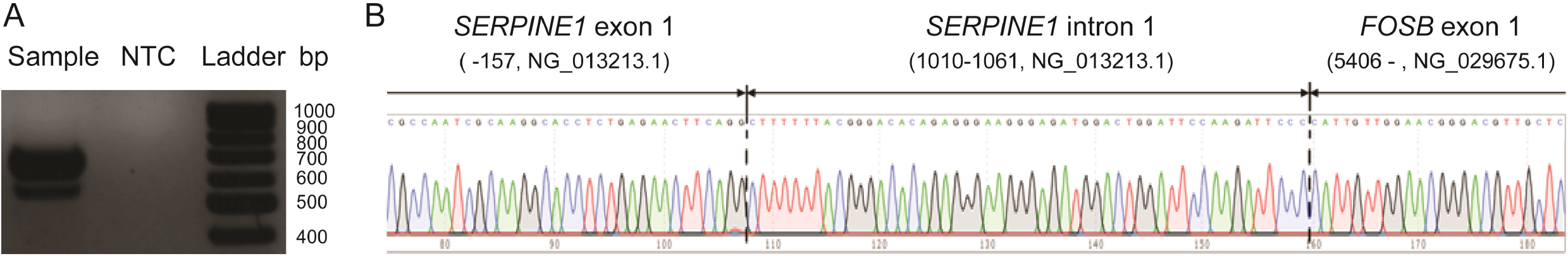
Identification of a *SERPINE1-FOSB* fusion gene. (A) Amplification of the fusion gene with a specific primer pair. (B) Sanger sequencing of the amplicon showing the structure of the fusion gene. NTC: no template control.

## Discussion

Genetic aberrations play important roles in the occurrence, development, and treatment of various human tumors. In PHE, chromosomal rearrangements leading to the fusion of *FOSB* (the whole or partial coding region) with other genes, such as *SERPINE1, ACTB, WWTR1*, and *CLTC*, have been identified.^2, 4-11^ The common feature of these cases is that the fusion introduces a strong promoter, resulting in deregulated overexpression of intact, partial, or chimeric FOSB. However, the pathogenic mechanism by which these genetic aberrations lead to PHE, which is of value for the development of effective targeted treatments, remains largely unknown. Recently, van IJzendoorn et al.^16^ demonstrated that a *SERPINE1-FOSB* fusion-derived chimeric protein could function as an active transcription factor that was not only capable of regulating its own transcription but also upregulating the expression of PDGFRA and FLT1. In their case, the tumors were responsive to telatinib, a drug currently in clinical trials, that can block self-regulation of the fusion gene by specifically affecting PDGFRA, FLT1, and FLT4 signaling and downregulating SERPINE1. However, the presence of different genetic aberrations, such as fusions with different genes or different domains, could alter the pathogenesis as well as the treatment. Therefore, further investigation is required to demonstrate whether this drug is effective in cases with other genetic aberrations such as *SERPINE1-FOSB* fusions with intact or truncated *FOSB* or *ACTB-FOSB* fusions. Similar to a previous case,^15^ in our case, mTOR was overexpressed as was pAKT (Fig. 3), suggesting activation of the PI3K-AKT-mTOR signaling pathway, one of the most frequently dysregulated pathways in human cancers that regulates important cellular functions, including metabolism, motility, growth, and proliferation.^17^ Dysregulation of this pathway supports the survival, expansion, and dissemination of cancer cells.^17^ Genetic alterations in specific oncogenes or tumor suppressor genes, such as *PIK3CA, PIK3R1, AKT, MTOR, TSC1*, and *TSC2*, can aberrantly activate this pathway via diverse mechanisms. However, these genetic alterations also provide opportunities for therapeutic targeting of this pathway and potential predictive biomarkers of therapeutic effectiveness.^17^ Currently, several distinct classes of drugs targeting the PI3K-AKT-mTOR signaling pathway, including PI3K and AKT inhibitors, as well as mTOR inhibitors, have been developed and have shown clinical potential.^18^ mTOR inhibitors, such as sirolimus and everolimus, have been effectively used for targeted therapy of PHE.^12-15^ However, the genetic information for mTOR inhibitor-responsive cases is limited. In some cases, the genetic information was missing,^14^ while in others, although a translocation or gene fusion was reported, the translational products of the fusion gene were not described.^13, 15^ Most recently, Bridge et al.^7^ reported a *CLTC*-*FOSB* fusion gene resulting in a chimeric protein in a case of PHE, and the tumor was responsive to sirolimus therapy. In our study, we clearly correlated the effectiveness of sirolimus therapy with a PHE-related genetic aberration, that is, a *SERPINE1*-*FOSB* fusion resulting in overexpression of intact FOSB. This genetic aberration could be a predictive biomarker for effective sirolimus therapy. FOSB belongs to the Fos family, which is associated with various cellular functions, including proliferation, differentiation, and transformation, and it is frequently involved in the pathogenesis of vascular tumors.^19-22^ However, the causal relationship between FOSB overexpression and activation of the PI3K-AKT-mTOR signaling pathway remains unclear. Nonetheless, the use of combinations of inhibitors targeting the PI3K-AKT-mTOR signaling pathway in PHE cases is worthwhile, since this strategy has been shown to be more effective than monotherapy, owing to the inhibition of compensatory mechanisms involved in intrinsic and adaptive resistance.^23, 24^ Future investigations of the interconnecting cascades between the PHE-related genetic aberrations and this signaling pathway will advance our knowledge of PHE and aid in the development of personalized agents with greater target specificity and optimal therapeutic index.

In conclusion, we report a novel *SERPINE1*-*FOSB* fusion gene in a patient with PHE that was responsive to sirolimus therapy. The fusion gene resulted in overexpression of intact FOSB, pAKT, and mTOR, suggesting activation of the PI3K-AKT-mTOR signaling pathway. Our study correlated the dysregulation of a specific signaling pathway and the effectiveness of a specific targeted therapy to a specific genetic aberration, which could promote the investigation of therapeutic targets and provide a predictive biomarker of therapeutic effectiveness for PHE with this type of genetic aberration.

## Acknowledgments

We thank Dr. Qinhe Fan and Fanfan Shu for their kind supports to this work. This work was supported by the Key Clinical Specialty Discipline Construction Program of Fujian Province (Grant No. 050145).

## Notes

### Competing Interest Statement

The authors have declared no competing interest.

